# A single DNA methylation site regulates cell fate during *Clostridioides difficile* sporulation

**DOI:** 10.64898/2025.12.24.696351

**Authors:** Pola Kuhn, John W. Ribis, Mi Ni, Gang Fang, Aimee Shen

## Abstract

DNA methylation is a widespread phenomenon in bacteria that can regulate gene expression, although the mechanisms underlying this epigenetic regulation are often poorly understood. In *Clostridioides difficile*, the orphan DNA methyltransferase CamA promotes sporulation, a process critical for the persistence and transmission of this nosocomial pathogen. However, the specific CamA target genes that drive this increased sporulation phenotype were unknown. Here, we show that methylation of a single CamA motif in the promoter region of *spoIIE*, which encodes a factor critical for activating the early-acting sporulation sigma factor σ^F^, is sufficient to promote *spoIIE* transcription, σ^F^ activation, and spore formation. Surprisingly, the CamA-dependent increase in *spoIIE* expression also increases the frequency of premature σ^F^ activation prior to asymmetric division, resulting in miscompartmentalization of σ^F^ activity. While this premature activation event triggers cell lysis in the well-studied spore-former *Bacillus subtilis*, we show that *C. difficile* cells retain developmental plasticity: predivisional cells that have prematurely activated σ^F^ can abort sporulation and resume vegetative growth, whereas cells that activate σ^F^ in the forespore after asymmetric division remain committed to sporulation. Thus, DNA methylation controls a critical cell fate decision in *C. difficile* without compromising its capacity to adapt to fluctuating environmental conditions. Finally, we show that CamA confers a significant fitness advantage during murine infection through mechanisms largely independent of its ability to promote sporulation. Since CamA is specific to *C. difficile* and epigenetically regulates multiple pathways critical for pathogen persistence, these analyses imply that CamA could be a promising antimicrobial target.

**Significance:** While DNA methylation of the bacterial genome is ubiquitous and can regulate diverse processes, identifying the methylation sites that drive a specific phenotype remains challenging. The *Clostridioides difficile*-specific orphan DNA methyltransferase CamA promotes sporulation and persistence in mice, but the mechanisms underlying these phenotypes were unknown. We identified a single methylation site in the promoter of a key sporulation gene that enhances sporulation, a process critical for this clinically relevant pathogen to transmit disease. Our analyses uncovered an unexpected developmental plasticity to *C. difficile*’s sporulation program while also revealing that CamA regulates processes beyond sporulation that are important for maintaining long-term infection. Thus, targeting CamA may offer therapeutic benefit.

## Introduction

While DNA methylation has historically been associated with restriction-modification systems that allow bacteria to defend against foreign DNA, the advent of methylome sequencing technologies has revealed that epigenetic gene regulation is widespread across the bacterial domain^1^. Indeed, over 93% of prokaryotic species encode at least one orphan DNA methyltransferase, which lack cognate restriction enzymes and can modulate diverse cellular processes^1–3^. Recent studies have implicated these enzymes in epigenetically regulating phenotypic heterogeneity, such as in the production of surface structures like LPS^4^, fimbriae^5^, and pili^6^. Although DNA orphan methyltransferases have been implicated in regulating diverse phenotypes, pinpointing the specific methylation sites that drive these effects remains challenging^7^. Given that DNA methylation can alter the 3D structure and function of the genome^8,9^, identifying the specific methylation sites that drive a specific phenotype is critical for distinguishing between global, pleiotropic effects and local, regulatory mechanisms.

We previously showed that the major nosocomial pathogen *Clostridioides difficile* encodes CamA, an orphan DNA methyltransferase that is strictly conserved across this genetically diverse species^10^. Since *C. difficile*’s core genome comprises less than 20% of its pan-genome, the conservation of *camA* implies that CamA-mediated DNA methylation plays an important role in regulating this organism’s physiology. Consistent with this hypothesis, we previously showed that CamA is a non-essential factor that promotes sporulation in *C. difficile*^10^, which is critical for the transmission of this obligate anaerobe^11^. We further found that loss of CamA impairs the ability of *C. difficile* to persist in a mouse infection model^10^. Given that spores contribute to *C. difficile*’s ability to persist in a host^12^, these two phenotypes may be interrelated. To gauge the extent to which the sporulation defect of a Δ*camA* mutant contributes to its persistence defect in the host, we sought to identify which of CamA’s nearly 8,000 recognition sites is responsible for promoting *C. difficile* sporulation.

Sporulation is a complex developmental program initiated in response to specific environmental cues, such as nutrient limitation^13^. The activation of the master transcriptional regulator Spo0A by these conditions leads to a transcriptional cascade that culminates in the formation of a metabolically dormant spore. Spo0A directly activates the expression of genes whose products mediate asymmetric division, the first morphological hallmark of sporulation^14,15^. The resulting mother and forespore cells have distinct fates due to the compartment-specific activation of four sporulation-specific sigma factors. The earliest acting sigma factor is σ^F^, and its activity is confined to the forespore despite σ^F^ being made throughout the predivisional cell^16,17^. Studies in *B. subtilis* have shown that σ^F^ is held inactive in the predivisional cell by a regulatory module consisting of SpoIIAA, SpoIIAB, and SpoIIE, which prevents its premature activation until after asymmetric division is complete^18–21^ (**Figure 1A**). This delay is critical for ensuring the compartment-specific activity of this sigma factor and the subsequent transcriptional cascade that allows the forespore to successfully differentiate into a mature spore. In *B. subtilis*, σ^F^ activation also commits a cell to sporulation^22^ such that the sporulating cell will complete its differentiation process regardless of whether nutrient-rich conditions are restored.

**Figure 1.**
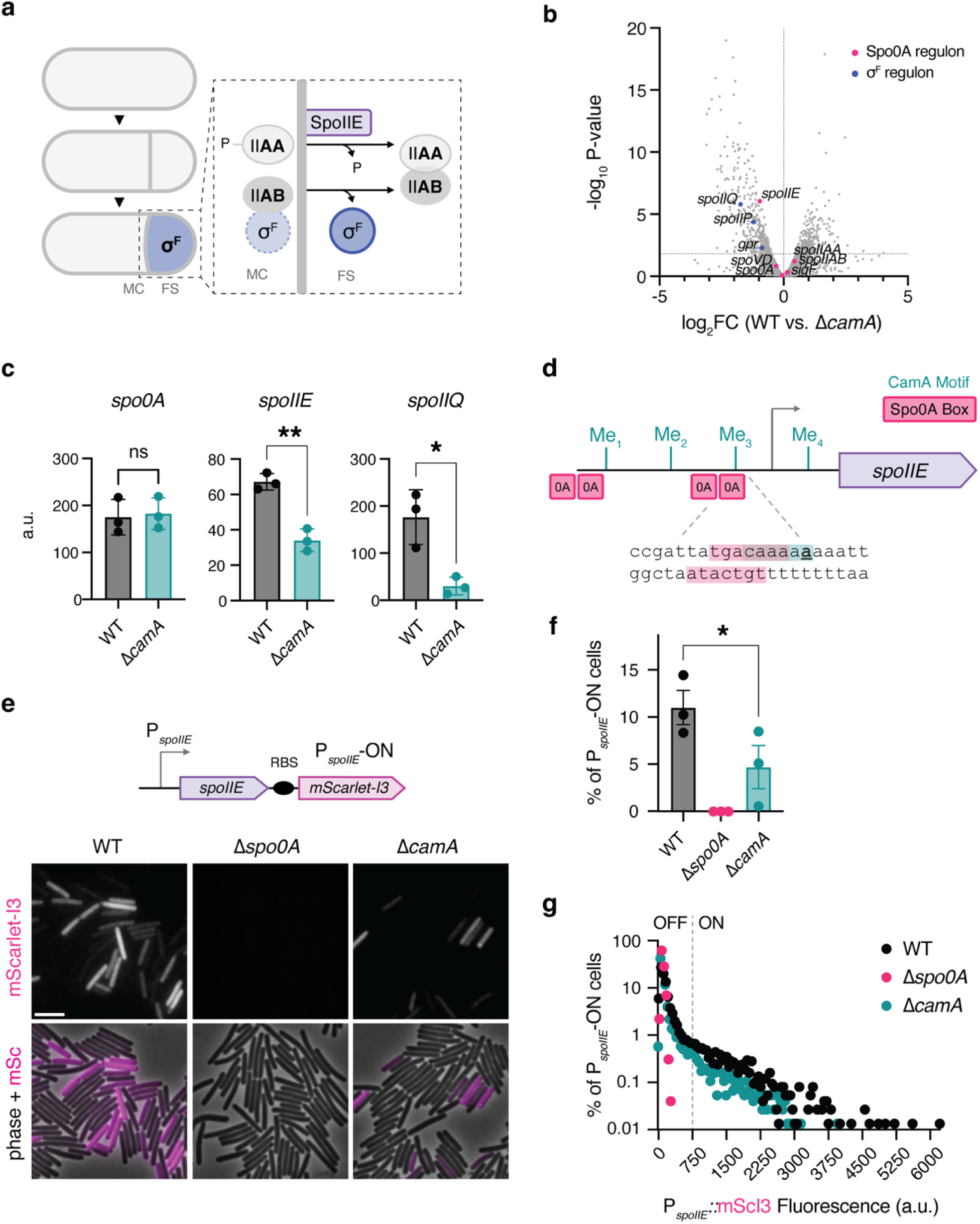
*spoIIE* is underexpressed in a Δ*camA* mutant unlike most Spo0A regulon genes. **(a)** Schematic of σ^F^ activation in *B. subtilis*. Before asymmetric division, σ^F^ is bound to and kept inactive by SpoIIAB. After asymmetric division, SpoIIE is preferentially localized to the forespore side of the polar septum where it dephosphorylates SpoIIAA, allowing it to sequester SpoIIAB. This releases σ^F^, enabling the transcription of the σ^F^ regulon in the forespore. **(b)** Volcano plot showing differentially expressed genes from RNA-seq analysis of WT and Δ*camA C. difficile* after 9 hours of growth on sporulation-inducing conditions (dataset was obtained from previous study^10^). The horizontal line indicates a p-value cutoff of 0.05. Spo0A regulon genes are labeled in pink, and σ^F^ regulon genes are labeled in blue. **(c)** Relative transcript levels as determined by RT-qPCR at 9 hours after sporulation induction. The data shown represent the mean ± standard deviation of three biological replicates from samples that were independent of those used for the RNA-Seq analyses. Statistical significance was determined using unpaired *t-*tests. *, p < 0.033; **, p < 0.002. **(d)** Schematic showing organization of the *spoIIE* promoter region in *C. difficile*. CamA motifs (CAAAA**<u>A</u>**) and Spo0A recognition sites (Spo0A box) are shown **(e)** Images of cells harboring a bicistronic *spoIIE-RBS-mScarlet-I3* transcriptional reporter construct integrated into the native *spoIIE* locus, grown on sporulation-inducing medium for 9 hours. Δ*spo0A* was used as a negative control, since this strain cannot initiate sporulation^64^. Scale bar, 5 µm. **(f)** Percentage of *spoIIE*-ON cells, as determined from quantified images of the strains shown in panel e. Statistical significance was determined using a one-way ANOVA and Tukey’s multiple comparisons test. *, p < 0.033. **(g)** Histogram showing the frequency distribution of cells at different levels of mean fluorescence intensity for strains shown in panel e (a.u. is arbitrary units). Data represent 7,500 cells counted across three biological replicates. Cells were classified as *spoIIE*-OFF or *spoIIE*-ON based on the bimodal distribution of mean fluorescence intensity observed in violin plots. The cutoff was set at the valley between the peaks and validated against fluorescence levels of cells barely detectable in microscopy images.

Using transcriptomic analyses, we previously showed that σ^F^ activation is the earliest sporulation event affected by the loss of CamA^10^, but it was unclear which aspect of the σ^F^ regulatory cascade is epigenetically regulated. Here, we identify a single methylation site in the promoter region of the critical sporulation gene *spoIIE* that promotes σ^F^ activation and increases *C. difficile* spore formation. Using single-cell analyses, we show that SpoIIE must accumulate to an optimal concentration to induce σ^F^ activation in the forespore. These analyses unexpectedly revealed that CamA-induced overexpression of *spoIIE* leads to the premature activation of σ^F^ in the predivisional cell in over a third of the sporulating population. Whereas this premature activation event can induce predivisional cells to lyse in *B. subtilis*^18,19,23^, we show that prematurely activating σ^F^ in predivisional *C. difficile* cells does not cause lysis or constrain them to completing sporulation. In contrast, *C. difficile* cells that activate σ^F^ in the forespore remain committed to completing their differentiation process even when favorable growth conditions return. Thus, our analyses reveal an unexpected flexibility in a critical developmental decision in *C. difficile* compared to *B. subtilis*. Finally, we show that the persistence defect of a Δ*camA* mutant is largely independent of its sporulation defect, indicating that CamA-mediated DNA methylation enhances the fitness of *C. difficile* by modulating processes beyond sporulation during infection.

## Results

### CamA promotes the expression of the early-acting and critical sporulation gene, *spoIIE*

Our previous RNA-Seq analyses indicated that the earliest sporulation stage altered in a Δ*camA* mutant relative to WT is the activation of σ^F^ because Spo0A regulon genes (like *spo0A* and the *sigF-spoIIAB-spoIIAB* operon) are similarly expressed in a Δ*camA* mutant relative to WT, whereas σ^F^ regulon genes (like *gpr*, *spoIIQ*, and *spoIIP*) are under-expressed^10^ (**Figure 1B**). Thus, to identify sporulation genes epigenetically regulated by CamA, we focused on genes involved in regulating σ^F^ activity. σ^F^ is constrained to the forespore even though it is produced throughout the predivisional cell. In *B. subtilis*, the compartment-specific activation of σ^F^ is controlled by a signaling cascade consisting of the anti-sigma-factor SpoIIAA, an anti-anti-sigma factor SpoIIAB, and the multi-functional SpoIIE phosphatase^18–21^. Since the genes encoding these regulatory factors are conserved in *C. difficile*^17^, we compared their expression levels using RT-qPCR. While the *sigF-spoIIAA-spoIIAB* operon was similarly transcribed in WT and Δ*camA*, *spoIIE* was under-expressed ∼2-fold in Δ*camA* relative to WT (**Figure 1C**), even though both the *sigF* operon and *spoIIE* gene are direct targets of Spo0A^13^. Intriguingly, the *spoIIE* promoter region is enriched in CamA recognition motifs (CAAAA<u>A</u>; the methylated base is underlined): the 160 bp region upstream of the *spoIIE* transcription start site (TSS) contains three methylation sites (**Figure 1D**). This represents a significant enrichment compared to equally sized regulatory regions (p < 10^-5^), and the presence of these sites is strictly conserved across 135 *C. difficile* genomes, despite the high degree of genetic variation between isolates (**Supplemental Figure 1**). Furthermore, the Me3 methylation site is directly adjacent to the predicted Spo0A binding site in the *spoIIE* promoter. Collectively, these analyses suggest an evolutionary pressure to maintain the CamA methylation sites in the *spoIIE* promoter region.

Although *spoIIE* expression was only reduced ∼2-fold in the bulk RNA-Seq and RT-qPCR analyses (**Figure 1B, C**), these population-wide measurements can obscure underlying cell-to-cell variability in transcription levels. Since DNA methylation can influence the proportion of cells exhibiting a given transcriptional state^9^, we hypothesized that CamA affects the frequency of *spoIIE*-expressing cells rather than the amplitude of gene expression. To test this, we generated a bicistronic *spoIIE*-RBS-*mScarlet-I3* transcriptional reporter by integrating an *mScarlet-I3* gene, carrying a *spoIIE* RBS, downstream of the *spoIIE* gene in its native locus. Analysis of *spoIIE* expression at the single-cell level after 9 hours of growth on sporulation-inducing medium revealed that ∼2-fold fewer Δ*camA* cells activated the P*_spoIIE_* transcriptional reporter compared to WT (4.7 ± 3.9% and 11.0 ± 3.1% of cells, respectively), (**Figure 1F**). The Δ*camA* mutant also induced *spoIIE* at lower levels, with the median fluorescence intensity of *spoIIE*-positive Δ*camA* cells being 1222 a.u. compared to 1311 a.u. for WT (**Figure 1G**). Together, these results suggest that CamA increases the frequency of cells that induce *spoIIE* expression, as well as the magnitude of gene expression at the single-cell level.

To determine the functional consequence of altering *spoIIE* expression levels in *C. difficile*, we first constructed and characterized a Δ*spoIIE* mutant. Similar to observations in *B. subtilis*, Δ*spoIIE* mutants failed to activate σ^F^ based on fluorescence microscopy analyses of a σ^F^-activity reporter (**Supplemental Figure 2A**) and did not form heat-resistant spores (**Supplemental Figure 2B**). In addition, Δ*spoIIE* cells failed to progress beyond asymmetric division and their polar septa were thicker relative to WT in transmission electron microscopy analyses (**Supplemental Figure 2C, D**), consistent with prior work in *B. subtilis*^24,25^. This indicates that *C. difficile* SpoIIE is essential for proper asymmetric division and σ^F^ activation; thus, it shares conserved functions with previously studied spore formers^25–27^. Taken together, these data indicate that CamA-mediated DNA methylation increases the proportion of cells expressing a gene that encodes a critical early-acting factor in the transcriptional program controlling sporulation.

### Methylation of a single site in the *spoIIE* promoter region enhances *C. difficile* sporulation levels

Since these analyses implied that CamA epigenetically regulates *spoIIE* expression to control *C. difficile* sporulation levels, we sought to directly test this hypothesis by determining the effect of mutating the CamA methylation sites in the *spoIIE* promoter region. To this end, we individually mutated the three methylation sites found upstream of the *spoIIE* TSS by first generating a deletion of this upstream region and then restoring the deleted region with the wild-type sequence (WT*) or sequences carrying mutations in the individual CamA recognition sites (CAAAA<u>A</u> ◊ CAAATA, Me1*, Me2*, and Me3*) (**Figure 2A**). Importantly, the mutations alter the CamA recognition sequence without altering the fifth adenine methylated by CamA, and the Me3* mutation does not affect the predicted Spo0A recognition motif close to the -35 site^28^. We then measured the sporulation efficiency of the resulting mutants using a heat resistance assay. While preventing the methylation of the Me1 or Me2 sites did not affect sporulation levels, preventing the methylation of the promoter-proximal CamA site (Me3*), which is adjacent to a predicted Spo0A binding site, phenocopied the sporulation defect observed in the Δ*camA* mutant (**Figure 2B**).

**Figure 2.**
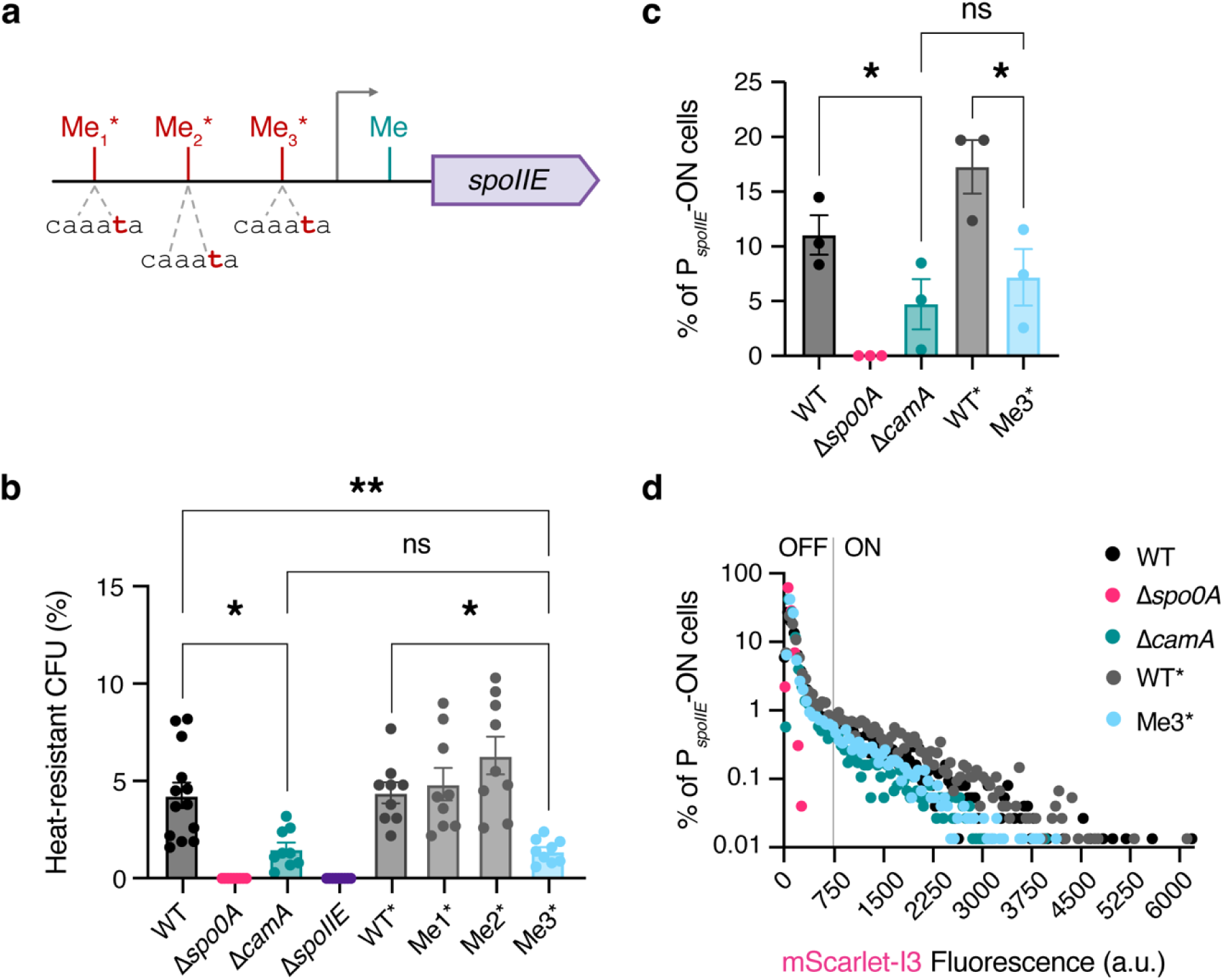
A single methylation site in the *spoIIE* promoter enhances sporulation. **(a)** Schematic of the individual methylation site mutations made in the *spoIIE* promoter region of *C. difficile* 630Δ*erm*. The region upstream of *spoIIE* was deleted, and DNA fragments containing the WT sequence with a watermark (WT*) or individual methylation site mutations (Me1*, Me2*, Me3*) were reintroduced by allelic exchange. **(b)** Heat resistance assay comparing the sporulation efficiencies of the methylation site mutants. Approximately 22 hours after sporulation was initiated, cells were heat-treated and then plated to enumerate the viable spore count relative to the untreated sample. Δ*spo0A* and Δ*spoIIE* were used as negative controls, since these strains cannot complete sporulation^64^. Results were analyzed using one-way ANOVA and Tukey’s multiple-comparisons test. *, p < 0.033. **, p < 0.002. Data represent a minimum of nine biological replicates. **(c)** Percentage of *spoIIE*-ON cells, determined using the threshold described above. Statistical significance was determined using a one-way ANOVA and Tukey’s multiple comparisons test. *, p < 0.033. **(d)** Histogram showing the frequency distribution of cells at different levels of mean fluorescence intensity for images of strains shown in Supplemental Figure 2. The data represent 7,500 cells counted across three biological replicates. Data for WT, Δ*spo0A* and Δ*camA* are also shown in Figure 1G and included here for direct comparison with WT* and Me3*.

To determine the effect of the Me3* methylation site mutation on *spoIIE* transcription, we introduced the bicistronic *spoIIE*-RBS-*mScarlet-I3* transcriptional reporter construct into the WT* and Me3* strains and analyzed *spoIIE* transcription using fluorescence microscopy. These analyses revealed that mutation of the single Me3 site was sufficient to phenocopy the decrease in *spoIIE* expression observed in Δ*camA* cells: fewer Me3* cells expressed *spoIIE* at detectable levels and the magnitude of their expression was reduced compared to WT* (**Figure 2C, D, Supplemental Figure 3**).

Since the Me3 motif partially overlaps with a putative Spo0A recognition motif (**Figure 1D**), we considered the possibility that mutation of the Me3 site affects the innate binding affinity of Spo0A for the P*_spoIIE_* promoter, even though the mutation is not predicted to alter the Spo0A recognition sequence^28,29^. Thus, we compared Spo0A binding to the Me3* and WT promoter regions using electromobility shift and fluorescence polarization assays. These assays both revealed that the DNA-binding domain of Spo0A binds to WT and Me3* P*_spoIIE_* DNA probes with similar affinity (**Supplemental Figure 4**). Thus, the reduced *spoIIE* expression and sporulation levels observed in Δ*camA* and Me3* mutants can be attributed to the loss of methylation, rather than a decrease in the intrinsic affinity of Spo0A for its binding sites. Together, these findings reveal that, of the nearly 8,000 CamA recognition motifs in *C. difficile*’s genome, methylation of the single Me3 site in *spoIIE*’s promoter region is sufficient to enhance *C. difficile* sporulation.

### Methylation of P*_spoIIE_* increases the frequency of σ^F^ activation in the forespore, as well as the predivisional cell

Since SpoIIE is predicted to be a positive regulator of σ^F^ activation (**Figure 1A**), we next asked whether the reduced *spoIIE* expression in the Δ*camA* and Me3* mutants is sufficient to reduce σ^F^ activation at the single-cell level. Analyses of σ^F^ activation using the σ^F^-dependent P*_gpr_::SNAP* transcriptional reporter (**Figure 3A**) after 15 hours of growth on sporulation-inducing medium revealed that 34% of WT cells activate σ^F^ in the forespore, while 17% of Δ*camA* and Me3* cells activate this sigma factor in this compartment (**Figure 3B, top panel**). These results are consistent with the ∼2-fold decrease in sporulation measured for cells that cannot methylate the Me3 site (Δ*camA* and Me3*, **Figure 2B**).

**Figure 3.**
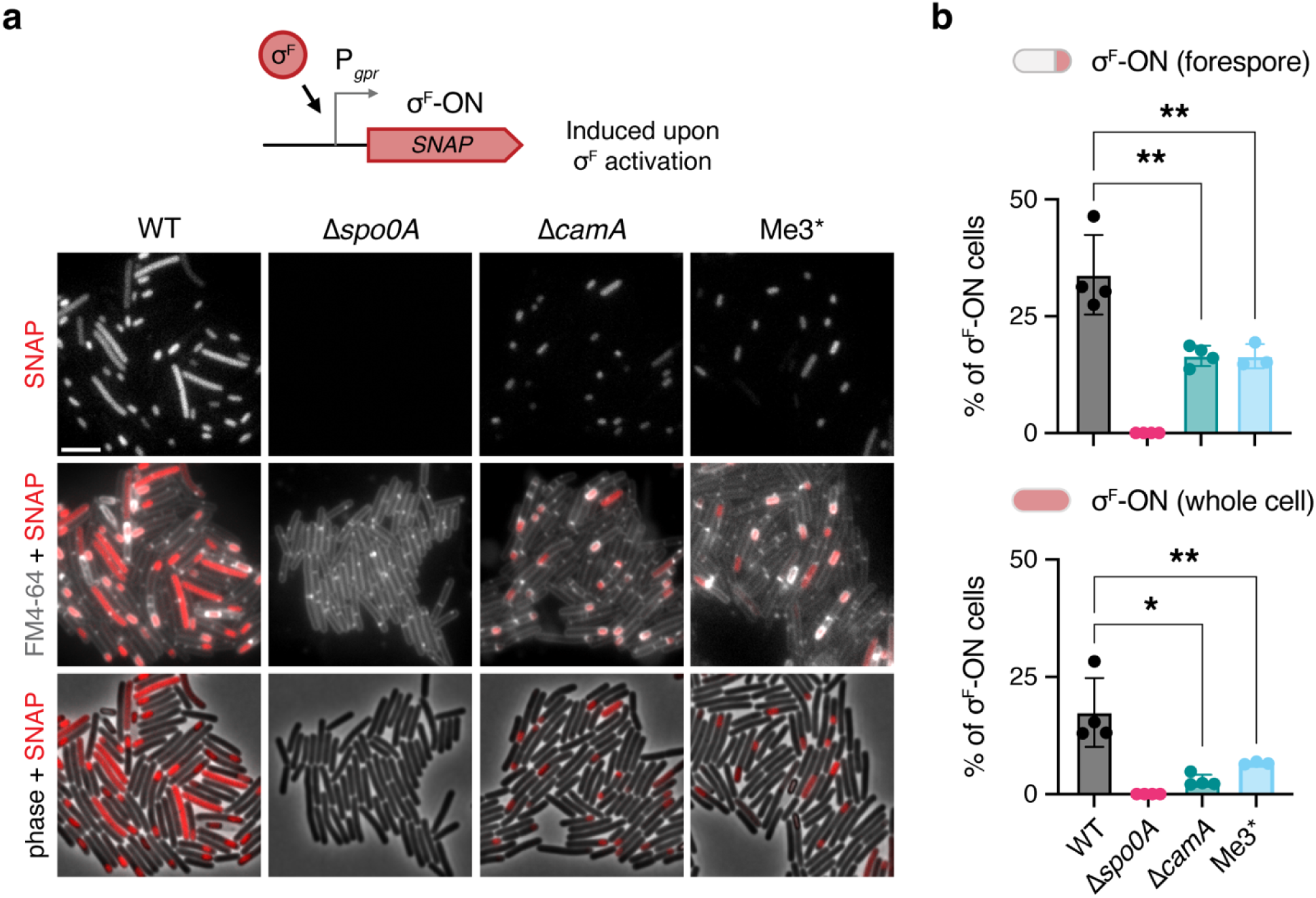
Methylation of P*_spoIIE_*enhances σ^F^ activation but impairs the compartmentalization of σ^F^ activity to the forespore. **(a)** Images of cells harboring a σ^F^ activity reporter construct (P*_gpr_::SNAP*) grown on sporulation-inducing medium for 15 hours. *gpr is* a member of the σ^F^ regulon, so its transcription serves as a reporter for σ^F^ activation. Cell membrane staining (FM 4-64) indicates that cells that have activated σ^F^ throughout the entire cell (labeled “whole cell”) have not completed asymmetric division. Δ*spo0A* was used as a negative control, since this strain cannot initiate sporulation^64^. Scale bar, 5 µm. **(b)** Percent of cells activating σ^F^ exclusively in the forespore or (**c**) throughout the whole cell for images shown in panel a. Data represent the mean ± standard deviation of at least 1,500 cells counted across three biological replicates. Statistical significance was determined using a one-way ANOVA and Tukey’s multiple-comparisons test. *, p < 0.033; **, p < 0.002.

Unexpectedly, the σ^F^ activity reporter revealed that a sizable proportion of visibly sporulating WT *C. difficile* cells activate σ^F^ throughout the entire cell, rather than exclusively in the forespore compartment. Specifically, 17% of WT cells activated σ^F^ in the entire cell compared to 3% and 7% of Δ*camA* and Me3* mutant cells, respectively (**Figure 3B, bottom panel**). Membrane staining revealed that cells with mislocalized σ^F^ activity throughout the cell have not yet completed asymmetric division, indicating that σ^F^ is being prematurely activated in predivisional cells (**Figure 3A**). In contrast, in *B. subtilis*, only 0.5-2% of sporulating cells activate σ^F^ in the predivisional cell^18,30–32^. Notably, in *B. subtilis*, the compartment-specific activation of σ^F^ in the forespore following asymmetric division is critical for the genetically identical mother and forespore cells to establish disparate transcriptional programs that allow for proper spore formation. Moreover, *B. subtilis* mutant cells that prematurely activate σ^F^ in the predivisional cell undergo lysis^18,19,23^, so the high proportion of σ^F^ activation in predivisional *C. difficile* cells was surprising.

Given the critical importance of properly compartmentalizing σ^F^ activity during *B. subtilis* sporulation, we sought to understand the basis and consequence of prematurely activating σ^F^ in predivisional *C. difficile* cells. We first considered whether the high frequency of premature σ^F^ activation was an artifact of the SNAP-tag labeling system. To test this, we visualized σ^F^ activation using a P*_spoIIQ_::mScarlet* transcriptional reporter, which is driven by a different σ^F^-dependent promoter and uses mScarlet as the visualizable reporter. With this system, we again found that 21% of WT cells activate σ^F^ in the predivisional cell (**Supplemental Figure 5**). Next, we wondered whether σ^F^ activation in the predivisional cells might be due to leakage of the reporter protein into the cytoplasm in cells that have retracted their polar septum, which can occur when the peptidoglycan synthesis or hydrolysis machinery is dysregulated during sporulation^33–35^. To assess this, we examined σ^F^ activation in Δ*spoIIP*Δ*spoIID* cells, which lack the septal hydrolysis machinery that mediates forespore cytoplasmic leakage in *B. subtilis*^33–35^. Whole cell σ^F^ activation was still observed in Δ*spoIIP*Δ*spoIID* cells (**Supplemental Figure 6**), indicating that the σ^F^ activity is indeed observed in predivisional cells rather than in cells that have retracted their septum. Taken together, these data indicate that σ^F^ activity is less tightly compartmentalized in *C. difficile* than in *B. subtilis* and that DNA methylation at the Me3 site of P*_spoIIE_* increases the frequency of premature σ^F^ activation in predivisional cells.

### Elevated *spoIIE* expression correlates with the loss of σ^F^ compartmentalization

Since sporulating WT cells express *spoIIE* at higher levels and activate σ^F^ prematurely in predivisional cells more frequently compared to the Δ*camA* and Me3* mutants (**Figure 3**), we hypothesized that elevated SpoIIE levels lead to the ectopic activation of σ^F^ prior to division. This hypothesis is based on observations in *B. subtilis*, where cells engineered to overexpress the *spoIIE* gene^18^ or accumulate the SpoIIE protein^26,36^ exhibit premature activation of this sigma factor. To test this possibility, we constructed a dual reporter system where (i) P*_spoIIE_* expression was detected using our bicistronic *spoIIE*-RBS-*mScarlet* transcriptional reporter integrated into the native locus and (ii) σ^F^ was monitored using a P*_gpr_::SNAP* transcriptional reporter construct integrated into the ectopic *pyrE* locus (**Figure 4A**). The induction of these reporters was analyzed after 12 hours of growth on sporulation-inducing medium, when both the *spoIIE* and σ^F^ activation reporters were visible. Indeed, these analyses revealed that, across all strains, forespore-specific activation of σ^F^ occurs within a relatively narrow range of *spoIIE* transcription (860 - 4680 a.u., median = 1946 a.u.), whereas σ^F^ activation in predivisional cells occurs at a higher range of *spoIIE* expression (1600 - 5447 a.u., median = 2971 a.u.) (**Figure 4B, Supplemental Figure 7**). These data strongly suggest that σ^F^ activation is sensitive to SpoIIE levels and that elevated levels of *spoIIE* transcription induces the premature activation of σ^F^ prior to the completion of asymmetric division in *C. difficile*, as has been documented in *B. subtilis* cells artificially engineered to overexpress *spoIIE*^18^. We next attempted to directly correlate SpoIIE levels and its localization to the polar septum with σ^F^ activation, by generating strains encoding SpoIIE-mScarlet-I3 protein fusions with different linkers. Unfortunately, the mScarlet-I3 variant was cleaved off all the fusions tested (data not shown), so we were unable to assess the relationship between SpoIIE localization and σ^F^ activation.

**Figure 4.**
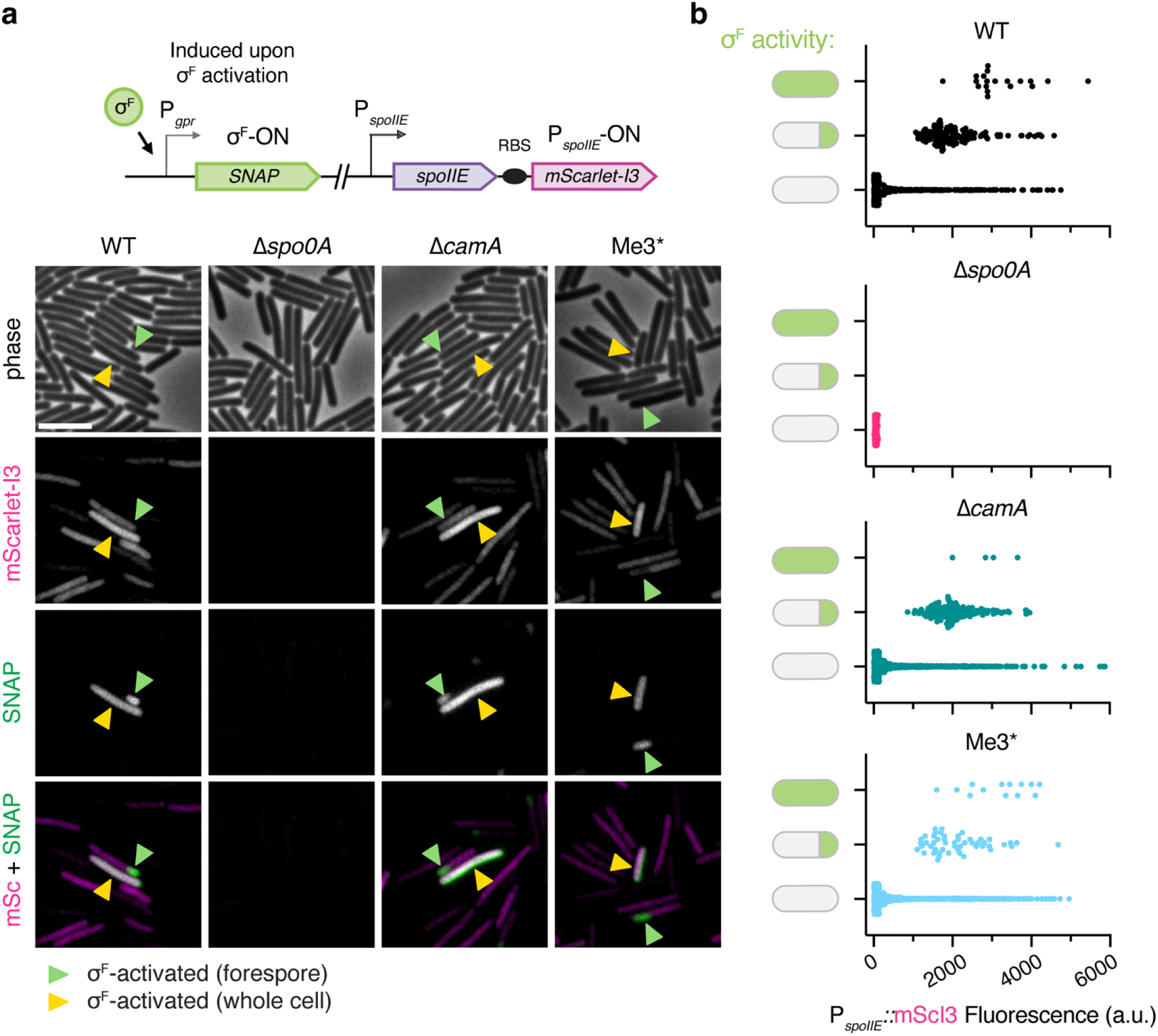
Elevated *spoIIE* expression correlates with the loss of σ^F^ compartmentalization. **(a)** Images of cells harboring a σ^F^ activity reporter (P*_gpr_::SNAP*) in the ectopic *pyrE* locus and a bicistronic *spoIIE-RBS-mScarlet-I3* transcriptional reporter integrated into the native *spoIIE* locus. Cells were grown on sporulation-inducing medium for 12 hours before labeling and fixation. Δ*spo0A* was used as a negative control, since this strain cannot initiate sporulation^64^. Green arrows indicate cells that have activated σ^F^ in the forespore; yellow arrows indicate cells that have activated σ^F^ across the whole cell. Scale bar, 5 µm. **(b)** Mean fluorescence intensities of cells grouped by their σ^F^ activation pattern (i.e., P*_gpr_::SNAP* expression): no σ^F^ activity, σ^F^ activity localized to the forespore, or σ^F^ activity observed across the whole cell. Due to day-to-day variability in the fluorescent signal from the *mScarlet-I3* reporter, the data shown are derived from a single biological replicate (1000 cells analyzed) that is representative of the trends observed for three independent replicates. The additional replicates and the resulting statistical analyses are shown in Supplemental Figure 6.

In *B. subtilis*, the coordinated regulation of SpoIIE’s localization, stability, and phosphatase activity tightly restricts σ^F^ activation to the forespore, even though all σ^F^ regulatory factors are present in the predivisional cell^18,20,37,38^. To probe how elevated SpoIIE levels promote σ^F^ activation in the predivisional cell, we first asked whether the systems that regulate SpoIIE in *B. subtilis* are conserved in *C. difficile*. In *B. subtilis*, the scaffolding protein DivIVA biases SpoIIE to the forespore face of the polar septum, which contributes to compartment-specific σ^F^ activation^30,31,39,40^. If similar DivIVA-dependent mechanisms regulate SpoIIE localization in *C. difficile*, deletion of *divIVA* should disrupt the architecture of septal proteins and impair σ^F^ compartmentalization. To test this hypothesis, we generated an in-frame deletion of *divIVA* and analyzed the activation of σ^F^ using the P*_gpr_::SNAP* reporter. In contrast with *B. subtilis*^31,41^, loss of DivIVA in *C. difficile* did not impact the ability to activate σ^F^ in the forespore (**Figure 5A**) or cause a sporulation defect. Instead, the sporulation efficiency of our Δ*divIVA* mutant was higher than WT, although this may be an artifact of the Δ*divIVA* chaining phenotype reducing the accuracy of cell counts and complicating the heat-resistant CFU measurements (**Figure 5B**). Complementation of the Δ*divIVA* strain with an ectopic copy of *divIVA* reversed the chaining and elevated sporulation phenotype (**Figure 5B**).

**Figure 5.**
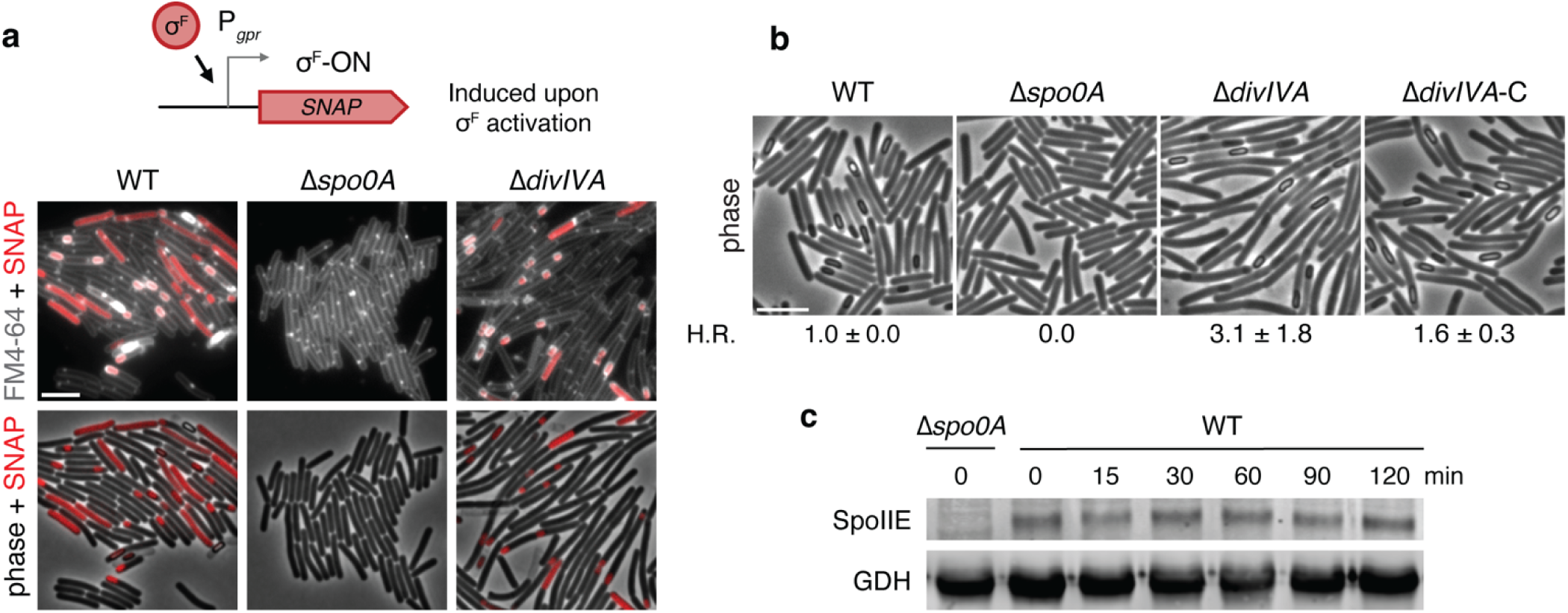
Forespore-specific σ^F^ activation in *C. difficile* is not dependent on canonical mechanisms for ensuring the compartmentalization of SpoIIE activity in *B. subtilis*. **(a, b)** Forespore-specific σ^F^ activity is independent of the structural protein DivIVA. **(a)** Representative images of cells harboring a σ^F^ activity reporter construct (P*_gpr_::SNAP*) in the *pyrE* locus grown on sporulation-inducing medium for 14-15 hours. Cell membrane staining (FM 4-64) indicates that the Δ*divIVA* mutant has a cell separation defect, but it can still compartmentalize σ^F^ activity to the forespore. Scale bar, 5 µm. **(b)** Phase-contrast images of WT and mutant strains grown on sporulation-inducing medium for 24-26 hours show that Δ*divIVA* cells sporulate efficiently. H.R. refers to the sporulation efficiency of mutants relative to WT based on their heat resistance properties. Sporulating cells were either heat-treated or left untreated prior to plating the samples on rich medium containing taurocholate germinant. The viable spore count was determined relative to the untreated sample; the resulting ratio was compared to the ratio determined for WT. Δ*spo0A* was used as a negative control, since this strain cannot initiate sporulation^64^. The data shown represent the mean ± standard deviation of at least three biological replicates. **(c)** Western blot analyses of SpoIIE stability following the arrest of transcription with rifampicin (100 µg/mL) and translation with chloramphenicol (100 µg/mL). Growth curves indicating *C. difficile*’s susceptibility to these antibiotics are shown in Supplemental Figure 7. Samples were collected at the indicated times after the addition of antibiotics and probed with anti-SpoIIE and anti-GDH (loading control) antibodies.

Another mechanism that promotes SpoIIE activity specifically in the forespore is its selective degradation by FtsH in the mother cell^30^. Notably, SpoIIE and FtsH are encoded next to each other in the genome of most spore formers, including *B. subtilis*, and a conserved N-terminal sequence in *B. subtilis* SpoIIE targets it for degradation^26,30^. However, *C. difficile spoIIE* lacks synteny with the *ftsH* gene and the N-terminal of *C. difficile* SpoIIE does not share conserved features with the FtsH-targeted degradation tag of *B. subtilis* (**Supplemental Figure 8**). Consistent with the absence of these features, we found that *C. difficile* SpoIIE remains stable for at least 120 minutes after both transcription and translation are arrested during sporulation (**Figure 5C, Supplemental Figure 9**). Taken together, these findings suggest that the forespore-specific activation of σ^F^ is not dependent on canonical mechanisms for compartmentalizing SpoIIE activity in *B. subtilis*, with mechanisms beyond SpoIIE stability or DivIVA-directed localization constraining *C. difficile* SpoIIE activity to the forespore.

### Activation of σ^F^ in the predivisional cell does not inhibit vegetative cell growth

In *B. subtilis*, the activation of σ^F^ is a critical developmental decision. If a sporulating culture is returned to nutrient-rich conditions, cells can revert to vegetative growth if they have not activated σ^F^ in the forespore. If they have activated σ^F^ in the forespore, the cell is irreversibly committed to completing this differentiation process regardless of the environmental conditions encountered^22,42^. Conversely, if a predivisional *B. subtilis* cell prematurely activates σ^F^, the cell will lyse due to mechanisms that remain poorly understood^18,19,23^. Given that σ^F^ is prematurely activated in a third of sporulating *C. difficile* cells (**Figure 3**), we wondered how this ectopic activation affects the fate of these cells and whether cells that have activated σ^F^ in the forespore are committed to completing sporulation, as has been documented in *B. subtilis*^22^. To address the latter question, we used time-lapse microscopy to study the fate of sporulating WT, *sigF* ^−^, and *sigE*^−^ cultures after they were returned to nutrient-rich conditions using an anaerobic imaging system we recently optimized for *C. difficile*^43^. Consistent with studies in *B. subtilis*, cells capable of activating σ^F^ (WT and *sigE*^−^) were unable to resume vegetative growth upon return to nutrient-rich conditions, provided that they had completed asymmetric division, whereas cells deficient for σ^F^ (*sigF*^−^) rapidly resumed vegetative growth despite having divided asymmetrically (**Supplemental Figure 10, Movies 1-3**). Thus, σ^F^ is critical for committing *C. difficile* cells to completing sporulation.

We next examined the consequence of prematurely activating σ^F^ in predivisional *C. difficile* cells using time-lapse microscopy. To identify sporulating cells that had activated σ^F^, we used WT and Δ*camA* strains harboring the σ^F^ activity reporter P*_gpr_::SNAP*. Sporulating cells that had activated σ^F^ were first labeled with the SNAP-tag reagent under anaerobic conditions^44^ and then inoculated onto nutrient-rich medium. These analyses revealed that predivisional cells that had prematurely activated σ^F^ throughout the cell were competent to resume vegetative growth, whereas cells that properly compartmentalized σ^F^ activity in the forespore were not (**Figure 6A, Movies 4-5**). To determine whether cells that activate σ^F^ in the predivisional cell were capable of activating σ^E^, the second sigma factor in the sporulation cascade, we introduced a σ^E^-dependent transcriptional reporter (*sipL*-RBS-*mScarlet-I3*) into a WT strain carrying the σ^F^-dependent transcriptional reporter (P*_gpr_::SNAP*). Analyses of this dual reporter strain revealed cells that activate σ^F^ in the forespore activate σ^E^ in the mother cell, while cells that prematurely activate σ^F^ in the predivisional cell fail to activate σ^E^, indicating they do not proceed with the sporulation program (**Figure 6B**). Taken together, our data reveal that, despite the irreversible nature of σ^F^ activation in the forespore (**Supplementary Figure 10**), σ^F^ activation in the predivisional cell permits exit from the sporulation program and resumption of vegetative growth. This developmental flexibility presumably enhances the ability of *C. difficile* cells to adapt to fluctuating environments. Our analyses further suggest that CamA-mediated increases in *spoIIE* expression increase the frequency of cells that activate σ^F^ in both the forespore and the predivisional cell, allowing CamA to promote sporulation without compromising a population’s developmental plasticity.

**Figure 6.**
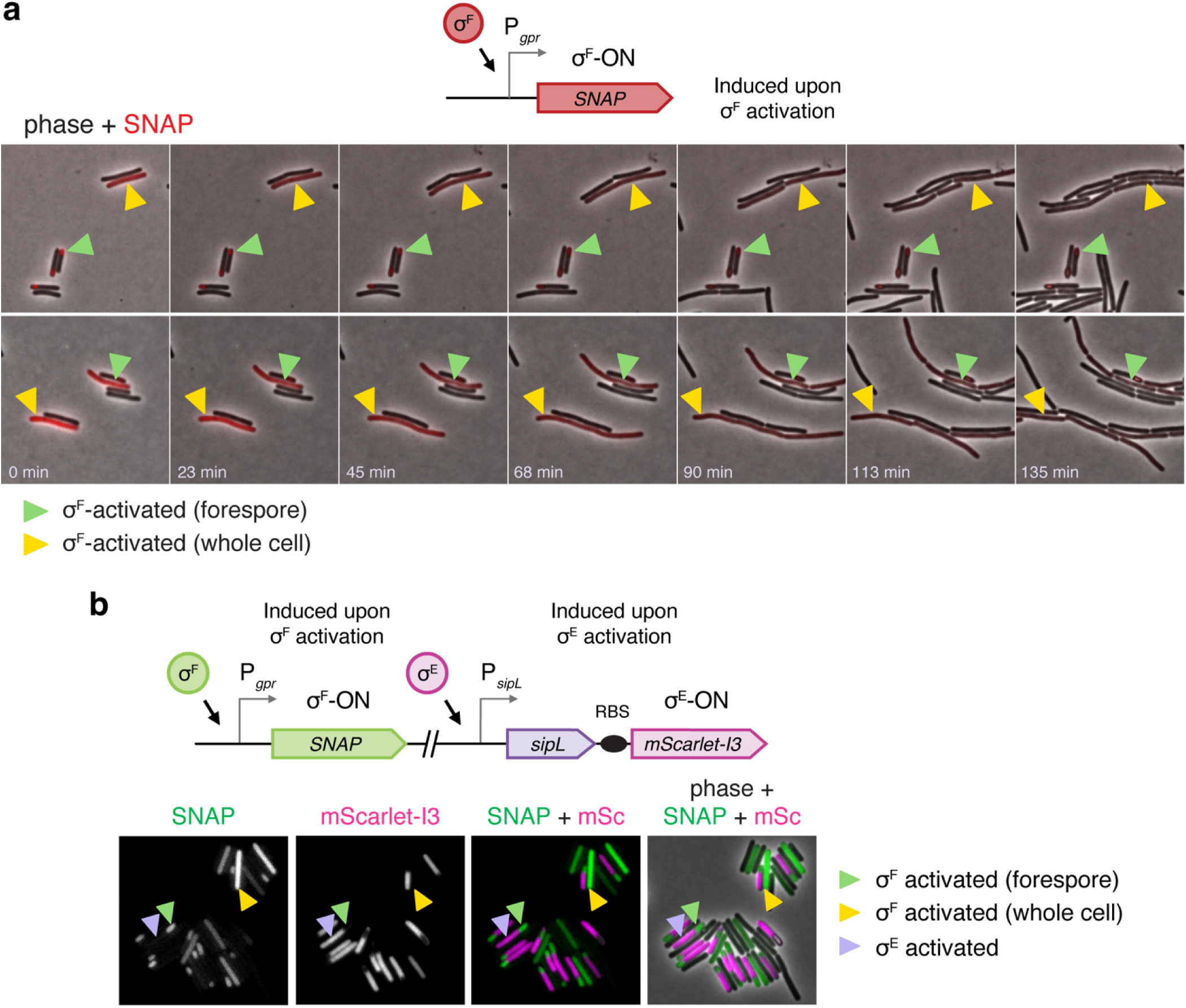
Cells that activate σ^F^ in the predivisional cell can resume vegetative growth. **(a)** Time-lapse microscopy analyzing the fate of sporulating WT cells that have activated σ^F^ in the forespore relative to the whole cell upon being transferred to nutrient-rich conditions. A WT strain carrying a σ^F^ activity transcriptional reporter (P*_gpr_::SNAP*) integrated into the *pyrE* locus was grown for 14 hours on sporulation-inducing medium. The cells were then labeled with SNAP-substrate to visualize cells that have activated σ^F^ and then spotted onto nutrient-rich agarose pads supplemented with cysteine. Agarose pads were prepared in the anaerobic chamber using gas-tight Gene Frames. After inoculating the agarose pad with the sporulating cultures, the growth chamber was sealed with a coverslip and then removed from the chamber for imaging. Cells were imaged every 2.5 min for approximately 4 hours. Green arrows indicate cells that have activated σ^F^ in the forespore; yellow arrows indicate cells that have activated σ^F^ across the whole cell. Corresponding movies are provided as Movies 4 and 5. Images are representative of three independent replicates. **(b)** Images of WT cells harboring a σ^F^ activity reporter (P*_gpr_::SNAP*) integrated into the ectopic *pyrE* locus and a bicistronic *sipL-RBS-mScarlet-I3* transcriptional reporter (reflecting σ^E^ activity) integrated into the native *sipL* locus. Cells were grown on sporulation-inducing media for 15 hours before labeling and fixation. Green arrows indicate cells that have activated σ^F^ in the forespore; yellow arrows indicate cells that have activated σ^F^ across the whole cell; purple arrows indicate cells that have activated σ^E^. Scale bar, 5 µm. Images are representative of three independent replicates.

### CamA confers a fitness advantage during infection that is largely independent of its effect on sporulation

Although the ability to sporulate does not influence *C. difficile*’s ability to cause gastrointestinal disease, it is essential for transmission between hosts^11^ and may contribute to persistence and disease recurrence^12^. Given the potential for mice to reinoculate themselves with spores through coprophagy, we sought to determine the extent to which Δ*camA*’s sporulation defect contributes to its persistence defect. To this end, we compared the relative fitness of the Me3* strain, which specifically recapitulates the sporulation defect of the Δ*camA* mutant, to WT and Δ*camA* in competition experiments. After sensitizing mice to infection with an antibiotic treatment regimen, mice were inoculated with a 1:1 mixture of spores consisting of pairwise combinations of WT, Δ*camA*, and Me3* (**Figure 7A**). To differentiate between strains, one member of each pair harbored a gene encoding spectinomycin resistance (*spec^R^*). To control for potential bias due to the presence of the *spec*^R^ cassette, we swapped the cassette between strains across replicates.

**Figure 7.**
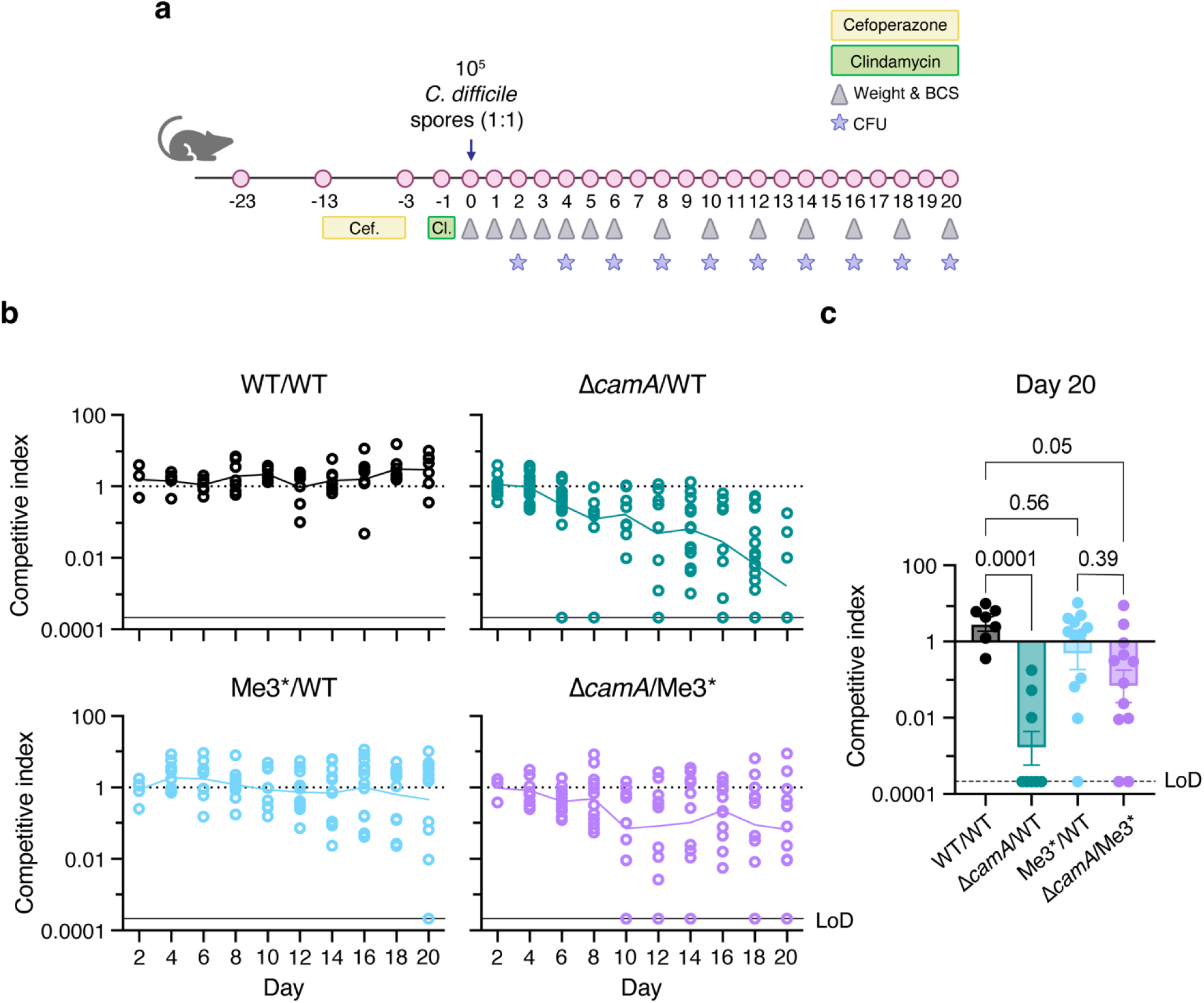
DNA methylation confers a multifactorial fitness advantage in a mouse model of infection. **(a)** Overview of the competitive infection model in mice. Female C57BL/6 mice were provided with water containing cefoperazone (0.5 mg/mL) for 10 days prior to an IP injection of clindamycin (10 mg/kg). Mice were inoculated via oral gavage with a 1:1 ratio of competing strains for a total dose of 10^5^ *C. difficile* spores. Bacterial CFU, mouse body weight, and body condition score were recorded at the indicated time points post-infection. The relative fitness of **(b)** WT/WT (2 experiments, 8 mice total), **(c)** Δ*camA*/WT (4 experiments, 18 mice total), **(d)** Me3*/WT (3 experiments, 12 mice total), and **(e)** Δ*camA*/Me3* (3 experiments, 12 mice total). The competitive index was calculated as (CFU_output, Strain A_/ CFU_output, Strain B_) ÷ (CFU_input, Strain A_/ CFU_input, Strain B_). **(f)** The competitive index of each pairwise combination on the final day of coinfection was determined and statistical significance was assessed using a one-way ANOVA and Tukey’s multiple-comparisons test.

As expected for infections with *C. difficile* 630Δ*erm*^45^, the mice exhibited mild weight loss and diarrhea at 2 days post-infection, but then regained the lost weight by Day 4. The mice remained stably colonized with *C. difficile* after their symptoms resolved and until the experiment was concluded (**Supplemental Figure 11)**. To validate our method of strain differentiation using spectinomycin resistance, we competed WT with the *spec*^R^-marked WT strain (WT/*spec*^R^); the strains exhibited equivalent fitness over the 20-day infection period (**Figure 7B**). However, by Day 14, WT had outcompeted Δ*camA* nearly to the limit of detection, and at the conclusion of the experiment, the mean competitive index (CI) was 0.0001 (**Figure 7C**). These data suggest that DNA methylation provides a fitness advantage for *C. difficile,* particularly during long-term colonization. To determine the extent to which Δ*camA*’s persistence defect is due to its impaired sporulation, we assessed the relative fitness of the Me3* mutant compared to WT and Δ*camA*. The Me3* mutant displayed a relatively mild fitness defect when co-infected with WT; this difference was not statistically significant (CI = 0.5) (**Figure 7C**). In contrast, Me3* outcompeted Δ*camA* (CI = 0.05) (**Figure 7C**). This suggests that the ability of CamA to enhance sporulation levels promotes *C. difficile*’s stable engraftment in a host, but it is not the primary driver of CamA’s fitness advantage. Together, these findings indicate that CamA-mediated DNA methylation regulates processes beyond sporulation that contribute to *C. difficile*’s ability to persist during murine infection.

## Discussion

Methylation of DNA by orphan DNA methyltransferases is widely observed in bacteria and has been shown to regulate gene expression and promote phenotypic heterogeneity in many bacterial systems^3,46–49^. However, beyond a few well-studied examples^3,9^, the regulatory mechanisms and functional consequences of these modifications are poorly understood. Here, we examined how CamA, an orphan DNA methyltransferase highly specific to the gastrointestinal pathogen *C. difficile*^10^, influences the differentiation from vegetative cells into endospores. Our analyses identified a single methylation site, among the 7,721 motifs methylated by CamA, that regulates sporulation in *C. difficile.* Specifically, we found that methylation of the *spoIIE* promoter region enhances the expression of *spoIIE*, increasing the frequency of σ^F^ activation and, thus, the proportion of *C. difficile* cells that form spores (**Figure 2**). Taken together, our study expands the relatively limited number of examples where the specific DNA methylation sites that regulate a given phenotype have been identified^7^.

Since we also demonstrated that σ^F^ activation in the forespore commits sporulating *C. difficile* cells to completing this differentiation process (**Supplemental Figure 10**), similar to *B. subtilis*^22^, our analyses reveal that *C. difficile* uses DNA methylation to regulate cell fate. Notably, the decision to sporulate is a high-stakes one: while spore formation is essential for *C. difficile* disease transmission^11^, committing to sporulation severely limits population growth, as a vegetative cell can produce millions of progeny during the time it takes to form a spore^42,50^. To remain adaptable, spore-forming bacteria take reversible steps towards an irreversible point of commitment^22,42^, up until which sporulating cells can revert to vegetative growth if environmental conditions improve. In *B. subtilis*, activation of σ^F^ in the forespore blocks the sporulating cell from reverting to vegetative growth^22^, while activation of σ^F^ in the whole cell appears to be lethal^18,19,23^.

In contrast, our study revealed an unexpected developmental flexibility to *C. difficile*’s sporulation process because we found that the activation of σ^F^ in whole cells does not cause developmental arrest or loss of viability in *C. difficile*. Instead, *C. difficile* cells that prematurely activate σ^F^ retain the capacity to resume growth when favorable conditions return (**Figure 6**). Since the ectopic activation of σ^F^ in predivisional cells is associated with high levels of *spoIIE* expression (**Figure 4**), our analyses reveal that DNA methylation saturates *spoIIE* expression to increase the proportion of *C. difficile* cells that form spores while also generating a subpopulation that remains adaptable to fluctuating environments.

Although we found that a specific methylation site regulates cell fate decisions in *C. difficile*, our study raises several questions. First, why can *C. difficile* abort the sporulation program after σ^F^ is activated in the predivisional cell and yet remain irreversibly committed to completing sporulation if σ^F^ is activated in the forespore? In *B. subtilis*, the expression of *spoIIQ* and *spoIIP* commits a sporulating cell to completing sporulation once σ^F^ has been activated^22^, but it remains unclear why predivisional cells that activate σ^F^ in *B. subtilis* induce cell lysis.

The precise mechanism by which DNA methylation of the Me3 site in the P*_spoIIE_* promoter region increases *spoIIE* expression is also unclear. Me3 overlaps a putative Spo0A recognition motif and is proximal to the -35 element (**Figure 1D**), so methylation may increase the binding affinity of Spo0A or RNA polymerase to this promoter. Notably, we did not detect appreciable differences in Spo0A binding affinity for methylated and unmethylated P*_spoIIE_* probes (**Supplemental Figure 12**). However, since these experiments were performed in the absence of RNA polymerase, they may not accurately recapitulate the transcriptional context found in *C. difficile*. Alternatively, it is possible that methylation of Me3 hinders the binding of a yet-unidentified transcriptional repressor, mimicking competitive binding models described in other systems^46,48,51^. In this scenario, a DNA-binding protein may compete with Spo0A for occupation of the Me3 site, hindering transcription in a proportion of the population.

Another key question is how *C. difficile* SpoIIE biases the activation of σ^F^ to the forespore. We found that two of the mechanisms that promote the compartment-specific activation of σ^F^ *in B. subtilis*, namely the preferential localization of *B. subtilis* SpoIIE to the forespore face of the polar septum by DivIVA^30,31,39,52^ and the targeted degradation of monomeric SpoIIE by FtsH in the mother cell, do not appear to regulate SpoIIE activity in *C. difficile*. Specifically, DivIVA does not affect the ability of *C. difficil*e to compartmentalize σ^F^ activity or complete the sporulation program, and *C. difficile* SpoIIE appears to be proteolytically stable (**Figure 5**). Consistent with this lack of post-translational regulation, our data suggest that the compartment-specific activation of σ^F^ occurs within a relatively narrow window of *spoIIE* transcription. If *spoIIE* expression exceeds this window, σ^F^ is more likely to be activated in the predivisional cell; conversely, underexpressing *spoIIE*, such as in the Δ*camA* and Me3* mutants, reduces the frequency of this ectopic activation event (**Figure 3**). We propose that, in the absence of multiple regulatory mechanisms controlling SpoIIE activity, *C. difficile* cells are more sensitive to elevated concentrations of this protein. Since multimerization of SpoIIE in *B. subtilis* controls its phosphatase activity^30,53,54^, *C. difficile* SpoIIE may be more likely to oligomerize in cells that express *spoIIE* at high levels in predivisional cells. Additional work is required to establish precisely how *C. difficile* compartmentalizes σ^F^ activity.

Finally, by demonstrating that the Me3* mutation recapitulates the sporulation defect of a Δ*camA* mutant, we were able to specifically assess how the CamA-mediated increase in sporulation contributes to the ability of *C. difficile* to persist within a murine host. Our finding that an Me3* mutant can persist in mice with similar efficiency as WT strongly suggests that CamA controls processes beyond sporulation that promote persistence in a host. Since the fitness advantage of WT relative to Δ*camA* was only observed approximately ten days post-challenge, DNA methylation would appear to be acting downstream of the acute infection phase. Importantly, the function of CamA has only been studied in a limited number of lab conditions, where it has been implicated in modulating biofilm formation, cell length, and flagellar gene expression^10^. Further investigation is required to establish how this enzyme promotes fitness during infection and whether CamA affects transmission or disease recurrence. Regardless, given its specificity to *C. difficile* and potential influence on multiple relevant pathways during infection, CamA may potentially serve as a viable therapeutic target. Taken together, this study underscores the capacity for DNA methylation to serve as a regulatory mechanism and provides a framework for mapping functional methylation sites in complex bacterial methylomes.

## Methods

### *C. difficile* strain construction and growth conditions

All *C. difficile* strains used in this study are listed in **Table S2** and are derivatives of 630Δ*erm*. Mutant strains were constructed in a 630Δ*erm*Δ*pyrE* strain using the *pyrE*-based allele-coupled exchange, as described previously^55^.

Strains were grown from frozen glycerol stocks on brain heart infusion medium supplemented with yeast extract (0.5% w/v) (BHIS) and taurocholate (0.1% w/v), thiamphenicol (10-15 µg/mL), kanamycin (50µg/mL), or cefoxitin (8 µg/mL) as needed. *C. difficile* defined medium (CDDM), supplemented with 5-fluoroorotic acid (2mg/mL) and uracil (5µg/mL) as needed, was used for constructing mutant strains. Sporulation analyses were carried out on cells cultured on 70:30 medium (70% BHIS and 30% SMC) as described previously^56^. For time-lapse experiments, strains were diluted into tryptone yeast extract broth supplemented with L-cysteine (0.1% w/v) (TYC) prior to inoculation onto 1.5% agarose pads supplemented with TYC medium. Resuspended and diluted fecal pellets collected during mouse experiments were plated on TCCFA plates with spectinomycin (500µg/mL) as needed. Cultures were grown at 37 °C under anaerobic conditions using a gas mixture containing 85% N_2_, 10% H_2_, and 5% CO_2_.

### *E. coli* strain construction and growth conditions

All *E. coli* strains used in this study are listed in **Table S3**, with links to plasmid maps containing primer sequences used for cloning. Plasmids were cloned using Gibson assembly and subsequently transformed into *E. coli* DH5ɑ. Isolated plasmids were sequence confirmed using Oxford Nanopore Technology before being transformed into *E. coli* HB101 for conjugation with *C. difficile* or *E. coli* BL21(DE3) for protein expression.

*E. coli* strains were grown in Luria-Bertani (LB) broth at 37 °C with 225 rpm shaking, supplemented with chloramphenicol (20 µg/mL), ampicillin (100 µg/mL), or kanamycin (30 µg/mL) as needed. For protein production, *E. coli* BL21(DE3) strains were grown in Terrific Broth (Thermo Fisher) supplemented with glycerol (0.5%), glucose (0.05%) and ɑ-lactose monohydrate (0.1%) at 20 °C for 60 hours with 225 rpm shaking.

### Sporulation induction

Starter cultures were grown in BHIS broth until they reached early stationary phase, then back-diluted to an OD_600_ of 0.05 and grown until they reached an OD_600_ of 0.35 to 0.75. 120 μL of these cultures were spread on 70:30 plates (40 mL media per plate).

### Heat resistance assay

After 22-24 hours of growth on 70:30 plates, sporulating cells were resuspended in phosphate-buffered saline (PBS), after which the culture was split into two. One sample was heat-treated at 60 °C for 30 minutes, while the other was plated. Both heat-treated and untreated samples were serially diluted and plated for viable count. Heat-resistance efficacies represent the average ratio of heat-resistant CFU to total CFU for a given strain relative to the ratio for the WT strain.

### Spore purification

After a minimum of 64 hours of growth on 70:30 plates, cells were resuspended in ice-cold sterile water, washed 5-8 times, and incubated on ice overnight. Samples were then treated with DNase I (New England Biolabs) for 1 hour at 37°C, washed an additional 2 times in ice-cold water, and purified with a 20% - 50% Histodenz (Sigma Aldrich) gradient. After 2 final washes with water, spore purity was assessed using phase-contrast microscopy, and cells were enumerated by viable count.

### RNA isolation

After 9-11 hours of growth on 70:30 plates, RNA was harvested from sporulating cells as previously described^57^. Briefly, samples were processed using the FastRNA ProBlue Kit (MP Biomedical) and a FastPrep automated homogenizer (three cycles of 40 seconds at setting 6.0; MP Biomedical). Contaminating genomic DNA was depleted by two sequential in-solution DNase treatments (New England Biolabs), followed by a column-bound DNase treatment with an RNeasy Kit (Qiagen). The DNase-treated RNA was enriched for mRNAs using a MICROB*Express* Bacterial mRNA Enrichment Kit (Invitrogen). Finally, the SuperScript First-Strand Synthesis System (Invitrogen) was used for reverse transcription of the enriched RNA to generate cDNA using random hexamer primers.

### RT-qPCR analysis

RT-qPCR was performed as previously described^10^, using the iTaq Universal SYBR Green supermix (BioRad) in a Mx3005P qPCR instrument (Stratagene). The following cycling conditions were used: 95°C for 2 minutes, 40 cycles of 95 °C for 15 seconds, and 60 °C for 1 minute. Transcript levels were normalized to the housekeeping gene *rpoB* using the standard curve method. Gene-specific primer pairs are listed in **Table S4**.

### Membrane, and SNAP labeling

FM4-64 (1 µg/mL; Invitrogen) was added to agarose pads to stain cell membranes. For labeling of strains encoding SNAP-tag fusions, cells were washed once with 0.5% Bovine Serum Albumin (BSA) in PBS and incubated with SNAP-tag 505-star or TMR-star (10 µM; New England Biolabs) at 37 °C in the dark. After 30 minutes, cells were washed three times in PBS and resuspended in PBS. All cell labeling was performed prior to fixation.

### Cell fixation

Cells were fixed as previously described^58^. Briefly, a 5X fixation solution (containing 20 µL 16% paraformaldehyde and 100 µL 1M NaPO_4_ buffer) was added to 500µL culture. Samples were incubated for 30 minutes at room temperature, followed by 30 minutes on ice. Fixed cells were washed three times in PBS and imaged within 48 hours of fixation.

### Microscope hardware

All samples were imaged on agarose pads made with TopVision Low Melting Point agarose diluted in PBS (1.5%; Thermo Fisher) and sealed with a coverslip. For phase-contrast and fluorescent micrographs, images were acquired using a Leica DMi8 inverted microscope equipped with a HC plan apochromat 64x1.5 NA oil immersion phase contrast objective, as previously described^43,59^. Excitation light was generated with a Lumencor Spectra-X multi-LED light source; for all fluorescent proteins aside for SNAP-tag 505-star, this was coupled with an XLED-QP quadruple-band dichroic beam splitter and an external emission filter wheel (Leica). FM4-64 was excited at 550/38 nm, and the emitted light was filtered using a 705/72 nm emission filter; mScarlet, mScarlet-I3 and SNAP-tag TMR-star were excited at 550/38 nm, and the emitted light was filtered using a 590/50 nm emission filter. Images of SNAP-tag 505-star were captured with a YFP filter set (Chroma), equipped with a 500/20 nm excitation filter, a 515-nm dichroic filter, and a 525/30 nm emission filter. 1 to 2 µm z-stacks were taken when needed. All imaging was carried out a 37 °C using a microscope incubation system (Pecon).

Phase-contrast microscopy without fluorescence was performed using a Zeiss Axioskop upright microscope with a 100x Plan-NEOFLAUR oil-immersion phase-contrast objective and a Hamamatsu C4742-95 Orca 100 CCD Camera.

### Time-Lapse Microscopy

Time-lapse imaging was performed as previously described^43^. Briefly, an anaerobic imaging chamber was constructed by layering two gas-impermeable 125 µL Gene Frames (Thermo Fisher) on a glass slide. Inside the anaerobic chamber, the Gene Frames were filled with pre-reduced TopVision Low Melting Point agarose (1.5%; Thermo Fisher) and TY media containing L-cysteine (0.1% w/v); a second glass slide was placed on the liquid media to create a flat surface; the slide was placed on a frozen freezer block until the agarose solidified. The second glass slide was removed, and the agarose pad was dried for 10 minutes prior to loading the cells and sealing the imaging chamber with a coverslip. The samples were imaged at 37° C at 2.5-minute intervals until they reached confluency in the field of view.

### Imaging analysis and quantification

After image acquisition, images were exported and processed in FIJI. Images were cropped to remove any out-of-focus cells, and the best-focused z-plane for each channel was selected to correct for chromatic aberration. To improve display, brightness and contrast settings were scaled and applied equally to all images shown in a single plane. At least three images were captured per replicate, and every strain was analyzed with three biological replicates. For cell segmentation and quantification of fluorescence intensities, images were additionally processed. First, Instant Computational Clearing (ICC) was performed (LASX software; Leica) to avoid bleed-through of fluorescent signal into neighboring cells. The adaptive strategy was run with the feature scale set to 2683 nm and 98% strength. Following ICC, images were processed as indicated above to remove all out-of-focus cells. To quantify the mean fluorescence intensity per cell, the MATLAB-based image analysis pipeline SuperSegger^60^ was used, with the supplied “60x *E.* coli” settings. For analysis of dual reporters presented in Figure 4, images were first processed via the SuperSegger image analysis pipeline, after which cells were manually categorized by σ^F^-activation status.

### Transmission Electron Microscopy

After 10 hours of growth on 70:30 plates, sporulating cells were fixed and processed for electron microscopy by the University of Vermont Microscopy Center as previously described^61^. Briefly, cells were fixed in 2% paraformaldehyde and 2% glutaraldehyde in 0.1 M sodium cacodylate buffer for 2 hours at 4°C, then washed with 0.1 M sodium cacodylate buffer. Samples were embedded in 2% agarose and cross-linked with 1% paraformaldehyde and 2.5% glutaraldehyde in 0.1 M cacodylate buffer, then washed with 0.1 M cacodylate buffer. After being minced into 1mm^3^ pieces, samples were dehydrated in a graded ethanol series (35%, 50%, 70% 85%, 95% and 100%) and cleared twice in 100% propylene oxide. Samples were infiltrated with Spurr’s epoxy resin in 100% polypropylene oxide in increasing ratios, then embedded in 100% Spurr’s resin and polymerized at 70°C. First, semi-thin sections (1 µm) were sliced on a Reichart Ultracut Microtome and stained with methylene blue-azure II; next, ultra-thin sections were cut with a diamond knife, retrieved on 200-mesh thin-bar nickel grids, and contrasted with uranyl acetate (2% in 50% ethanol) and Reynolds’ lead citrate. Images were captured on a JEOL 1400 Transmission Electron Microscope (Jeol USA).

### Protein purification

*E. coli* BL21(DE3) encoding lactose-inducible, His_6_-tagged proteins of interest (SpoIIE_Δ1-905_, Spo0A DNA binding domain (DBD), full-length Spo0A and CamA) were grown to stationary phase, back-diluted 1:1000 in 1 L Terrific Broth (Thermo Fisher) supplemented with glycerol (0.5%), glucose (0.05%) and ɑ-lactose monohydrate (0.1%). Cell cultures were grown at 20 °C with 225 rpm shaking; after 60 hours, cultures were pelleted, resuspended in 25 mL low imidazole buffer (LIB; 500 mM NaCl, 50 mM TRIS-HCl pH 7.5, 15 mM imidazole, 10% glycerol, 2 mM β-mercaptoethanol), and flash frozen in liquid nitrogen. Once thawed, cells underwent three cycles of probe sonication consisting of 45 seconds at 40% amplitude followed by 5 minutes on ice. Samples were pelleted at 1000 rpm for 45 minutes at 4 °C, and tagged proteins were affinity-purified from cleared lysates using Ni-NTA agarose beads with gentle rocking at 4 °C for 2 hours. Beads were washed three times with LIB and protein was eluted with high imidazole buffer (HIB; 500 mM NaCl, 50 mM TRIS-HCl pH 7.5, 200 mM imidazole, 10% glycerol, 2 mM β-mercaptoethanol). Beads were washed 6 times with HIB, pelleted, and the supernatant containing the eluted protein was collected and analyzed using Coomassie staining.

Affinity-purified proteins were then concentrated using an Amicon Ultra-15 10 kDa cutoff centrifugal filter (Millipore Sigma) and further purified by size exclusion chromatography (SEC) using a Superdex 200 Increase 10/300 GL column and 200 mM NaCl, 10 mM Tris pH 7.5, and on an AKTA pure protein liquid chromatography instrument. Fractions were collected every 0.5 mL, and fractions of interest were reconcentrated, aliquoted, and flash-frozen in liquid nitrogen.

### Fluorescence polarization assay

25-30 bp double-stranded DNA (dsDNA) was generated by incubating equal molar ratios of 6-Carboxyfluorescein-labeled oligonucleotides with their unlabeled complementary counterpart in annealing buffer (10 mM TRIS-HCL pH 7.5, 1 mM EDTA, 50 mM NaCl) at 95 °C for 15 minutes and gradually cooling the solution to room temperature. All oligonucleotide sequences are provided in **Table S4**. 1 nM dsDNA was mixed with serially diluted, purified Spo0A in fresh binding buffer (50 mM Tris-HCl pH 8, 100 mM KCl, 2.5 mM MgCl_2_, 0.2 mM DTT, 10% glycerol, and 2 µg salmon sperm DNA in H_2_O). After 10 minutes of incubation at room temperature, fluorescence polarization was read using a Synergy H1 plate reader (Agilent BioTek). 6-Carboxyfluorescein was excited at 485/20 nm, and emission was detected at 528/20 nm.

### Electromobility shift assays

250 bp DNA fragments encompassing the *spoIIE* promoter were amplified from purified *C. difficile* 630Δ*erm* genomic DNA using IRDye800-conjugated or regular primers (Integrated DNA Technologies). P*_spoIIE_*-specific pairs are provided in **Table S4**. The resulting labeled and cold competitor DNA probes were PCR- and gel-purified prior to use. 20 fmol labeled DNA (or 20 fmol labeled with 1000 fmol cold competitor DNA) was mixed with purified Spo0A-DBD-His_6_ in binding buffer (10 mM Tris-HCL, pH 7.6, 1 mM EDTA, 50 mM NaCl, 1 mM DTT, 5% glycerol) for 30 minutes at 37 °C. 6X loading dye (40% w/v sucrose, 0.25% bromophenol blue, and 0.25% xylene cyanol FF) was added to 1X before samples were loaded and run on an 8% native polyacrylamide gel at 80 V at 4 °C in the dark.

### Methylation of P*_spoIIE_*

The enzymatic activity of purified CamA-His_6_ was confirmed using the MTase-Glo^TM^ Methyltransferase Assay Kit^62,63^ (Promega). Briefly, 60 bp dsDNA were generated as described in the above section; all oligonucleotide sequences are provided in **Table S4**. In white assay plates (CORNING), 5 µM dsDNA was added to a mixture of 40 µM S-adenosyl-_L_-methionine and CamA at the indicated concentrations in a reaction buffer (80 mM Tris-Buffer, pH 8.0, 200 mM NaCl, 4 mM EDTA, 12 mM MgCl_2_, 0.4 mg/mL BSA, 4 mM DTT). After 30 minutes of incubation at room temperature, the protocol was completed per the manufacturer’s instructions. Luminescence signal was measured by a Synergy H1 plate reader (Agilent BioTek). To methylate DNA probes for electromobility shift assays, 200 nM DNA fragments, encompassing the *spoIIE* promoter, were incubated with 1 µM affinity- and SEC-purified CamA-His_6_ for 30 minutes at room temperature and PCR purified.

### Protein Degradation and Western Blot

After 11 hours of growth on 70:30 media, sporulating cells were resuspended in BHIS broth. Translation was inhibited with the addition of chloramphenicol (100 µg/mL) and transcription was inhibited by the addition of rifampicin (100 µg/mL); samples were removed at the indicated time points, pelleted at 4°C, and frozen at -80 °C. Samples were prepared for immunoblotting as previously described^57^. Briefly, they underwent three freeze-thaw cycles followed by the addition of EBB buffer (9 M urea, 2 M thiourea, 4% SDS, 2 mM beta-mercaptoethanol) and then were incubated at 95 °C with periodic vortexing. Pellets were resuspended, and bromophenol blue was added to 0.01% (w/v) to visualize the samples. Samples were vortexed vigorously, boiled, and pelleted again, immediately prior to loading on a gel. Proteins were resolved on a 12% SDS-polyacrylamide gel electrophoresis (SDS-PAGE) gel.

Proteins were transferred to polyvinylidene fluoride membranes (PVFD), which was washed and blocked with Odyssey blocking buffer (LiCor) for 30 minutes. PVFD membranes were then probed with rabbit anti-SpoIIE (1:1000 dilution) and chicken anti-GDH (1:5000 dilution; Thermo Fisher) polyclonal primary antibodies. After washing in PBS with Tween, PVFD membranes were probed with goat anti-rabbit IRDye680 and donkey anti-chicken IRDye800 secondary antibody (1:12,000 dilution; LiCor Biosciences).

### Growth Curve assays

Starter cultures were grown in BHIS broth until they reached early stationary phase and then back-diluted to an OD_600_ of 0.05. Once grown to mid-log phase, cultures were normalized to a starting OD_600_ of 0.5. Cells were diluted 1:30 in BHIS, with antibiotics supplemented as indicated, and 150 µL was distributed into wells in technical triplicate. Plates were incubated in an Epoch plate reader (Agilent BioTek) at 37°C in the anaerobic chamber with linear shaking every 2 minutes. OD_600_ values were recorded every 15 minutes.

### Mouse infection experiment

Groups of seven-week-old female C57BL/6 mice (purchased from Jackson Laboratory) were housed together in a large, sterile rat cage to normalize their microbiota. After 10 days, cefoperazone (0.5mg/mL) was administered in their drinking water, which was provided *ad libitum* for 10 days and replaced every 2 days. Mice were returned to regular drinking water for 2 days before receiving a single intraperitoneal injection of clindamycin (10 mg/kg in 200 μL PBS). After 24 hours, groups of 4 mice were transferred to smaller cages and inoculated via oral gavage with a total of 1 × 10^5^ purified spores in 200 µL PBS. The spore solution contained a 1:1 mixture of strains of interest, with one strain harboring the *aad9* gene encoding spectinomycin resistance integrated into the *pyrE* locus. Weight and body condition scores were recorded, and fecal pellets were collected at the indicated times. Fecal pellets were weighed prior to being resuspended in 1 mL PBS, and 10-fold dilutions were plated on TCCFA agar in the presence and absence of spectinomycin (500 µg/mL). Colony-forming units were counted and normalized to the mass of the original fecal pellet. The competitive index was calculated as (Output_SpecS_/Output_SpecR_)÷ (Input_SpecS_/Input_SpecR_), or vice versa. Cage changes were performed every 2 days, and mice were fed irradiated Lab Diet 2918 throughout. At the experimental endpoint, mice were euthanized with CO_2_ asphyxiation followed by cervical dislocation.

### Data visualization and statistics

EMSAs and western blots were visualized using a LiCOR Odyssey CLx Imager. All graphs and subsequent statistical tests were generated using GraphPad Prism.

## Notes

### Competing Interest Statement

The authors have declared no competing interest.

